# Frontal cortical functional connectivity is impacted by anaesthesia in macaques

**DOI:** 10.1101/2021.06.09.447698

**Authors:** C Giacometti, A Dureux, D Autran-Clavagnier, C. R. E. Wilson, J Sallet, M. Dirheimer, E Procyk, F Hadj-Bouziane, C Amiez

## Abstract

A critical aspect of neuroscience is to establish whether and how brain networks evolved across primates. To date, most comparative studies have used resting-state functional Magnetic Resonance Imaging (rs-fMRI) in anaesthetized non-human primates and in awake humans. However, anaesthesia strongly affects rs-fMRI signals. The present study investigated the impact of the awareness state (anaesthesia vs. awake) within the same group of macaque monkeys on the rs-fMRI functional connectivity (FC) organization of a well characterized network in the human brain, the cingulo-frontal lateral network. Results in awake macaques revealed a similar FC pattern to that previously uncovered in the human brain. Rostral seeds in the cingulate sulcus exhibited stronger correlation strength with rostral compared to caudal lateral frontal cortical areas while caudal seeds in the cingulate sulcus displayed stronger correlation strength with caudal compared to anterior lateral frontal cortical areas. Critically, this inverse rostro-caudal functional gradient was abolished under anaesthesia. This study demonstrates that the FC pattern of cingulo-frontal cortical networks is preserved from macaque to human but some of its properties can only be observed in the awake state, warranting significant caution when comparing FC patterns across species under different states.

## Introduction

A critical aspect of neuroscience is to identify whether and how brain networks evolved across primates in order to 1) establish the putative uniqueness of the human brain and 2) to allow an optimal transfer of results obtained in non-human primates (NHP) to humans.

Most comparative studies in the past decade have used resting-state functional Magnetic Resonance Imaging (rs-fMRI), a non-invasive approach focusing on the assessment of spontaneous low-frequency fluctuations of Blood-Oxygenation-Level-Dependent (BOLD) signal (<0.1Hz) at rest (Biswal et al., 1997). This method reveals temporal correlations of activity between brain areas (Biswal et al., 2010) and allow to compare brain anatomo-functional connectivity (FC) organization across primate species (Amiez et al., 2021; Barron et al., 2021; Folloni et al., 2019; Friedrich et al., 2021; Hadj-Bouziane et al., 2014; Hutchison et al., 2011, 2012, 2013, 2015; Lopez-Persem et al., 2020; Mars et al., 2011, 2013, 2016, 2018; Hutchison & Everling, 2012; Neubert et al., 2015, 2014; Sallet et al., 2013; Thomas et al., 2021; Van Essen et al., 2016; Vincent et al., 2007; Yin et al., 2019). For example, it has been shown that large-scale resting state networks (e.g. the default-mode network) are topographically and functionally comparable between humans and macaques (Hutchison et al., 2011; see also Mars et al. 2012). Although these rs-fMRI studies have provided evidence of functional homologies between human and non-human-primates, a critical issue is that they were performed in anaesthetized macaques and awake humans (PRIME-DE https://fcon_1000.projects.nitrc.org/indi/indiPRIME.html;Amiez et al. 2021; Folloni et al., 2019; Hutchison et al., 2012, 2013, 2015; Lopez-Persem et al., 2020; Neubert et al., 2014, 2015; Sallet et al. 2013; Thomas et al., 2021; Vincent et al., 2007; Yin et al., 2019). There is a growing literature showing that anesthetic drugs alter brain FC patterns both in humans (for review, see Alkire & Miller, 2005; Hudetz, 2012) and non-human primates (Areshenkoff et al., 2021; Barttfelda et al., 2015; Hori et al., 2020; Hutchison et al., 2014; Li et al., 2013; Lv et al., 2016; Rao et al., 2017; Signorelli et al., 2021; Uhrig et al., 2018; Wu et al., 2016). In the case of isoflurane, one of the most commonly used anesthetic agents, several major impacts on brain function have been described: 1) a global FC (i.e. correlation of activity between brain regions) breakdown when using a concentration in the inhaled air >1.5%, 2) a decrease of anticorrelations, 3) a stronger alteration of interhemispheric compared to intrahemispheric FC (Hutchison et al., 2014; Barttfeld et al., 2015; Hutchison et al., 2014; Hori et al., 2020; Uhrig et al., 2018; Xu et al., 2018).

Here we sought to determine the extent to which anaesthesia affects frontal cortical network connectivity. To do so, we identified in the same group of macaques, the impact of the awareness state (anaesthesia vs. awake) on the rs-fMRI FC organization of a well characterized network in the human brain, the cingulo-frontal lateral network (Loh et al. 2018), for which potential homologies in macaque have been described (for review see Procyk et al., 2016). In awake humans, three cingulate motor zones occupy the cingulate cortex along a rostro-caudal axis: the most rostral one is RCZa (anterior Rostral Cingulate Zone), RCZp (posterior Rostral Cingulate Zone) is observed caudally to RCZa, and finally CCZ (Caudal Cingulate Zone) occupies the most caudal location. RCZa and RCZp/CCZ have been shown to display a linear inverse FC gradient with the lateral prefrontal cortex and motor regions. RCZa displays stronger positive correlations with rostro-lateral prefrontal areas (e.g. frontopolar cortical area 10, Broca area (BA), dorsolateral prefrontal cortical areas 46 and 9/46) and weaker ones with the caudal lateral-frontal motor areas (e.g. Frontal Eye Field -FEF-, primary tongue and primary hand motor areas, called M1Face and M1Hand respectively). Conversely, RCZp and CCZ display the opposite pattern, i.e. weaker correlation strength with more rostral lateral-prefrontal areas and stronger correlation strength with more caudal lateral motor frontal areas (Loh et al., 2018). Macaque cingulate motor areas, named CMAr (rostral Cingulate Motor Area), CMAv (ventral Cingulate Motor Area), and CMAd (dorsal Cingulate Motor Area) have been suggested to be homologous to the human RCZa, RCZp, and CCZ, respectively (Amiez & Petrides, 2014; Picard and Strick 2001; Procyk et al. 2016). Furthermore, macaque brains do present homologues of a set of rostro-lateral frontal areas studied in humans (i.e. area 10, area 46, area 9/46, BA, FEF, M1Face and M1Hand areas; Petrides and Pandya, 1994). Based on these findings, we tested whether, in macaques, 1) the FC within the cingulo-frontal lateral network would follow a similar rostro-caudal functional gradient to that uncovered in the human brain and 2) this functional organization would be impacted by anaesthesia.

Three *rhesus* macaques underwent rs-fMRI sessions in both anaesthetized and awake states. Results revealed that 1) the inverse functional gradient displayed by rostral cingulate *versus* caudal cingulate regions with rostro-lateral prefrontal regions and caudal lateral motor frontal regions is highly preserved from awake macaques to humans and 2) it is abolished in anaesthetized compared to awake state in macaques.

## Results

We compared the FC pattern within the cingulo-frontal lateral network of three *rhesus* macaques under two states, awake and anaesthetized, using rs-fMRI. As summarized in Table 1, for each animal, we acquired six runs under anaesthesia (1-1.5% isoflurane) and 12 runs while the animals were awake. Each run comprised 400 volumes (see Methods). We assessed FC (Pearson correlation strengths) between 11 seeds located along the cingulate sulcus and 7 Regions Of Interests (ROIs) along a rostro-caudal axis in the lateral frontal cortex. Seeds were positioned in an antero-posterior extent that would fairly encompass all cingulate motor areas. ROIs were selected based on the known homologies between humans and macaques (Petrides and Pandya, 1994).

**Table 1.**
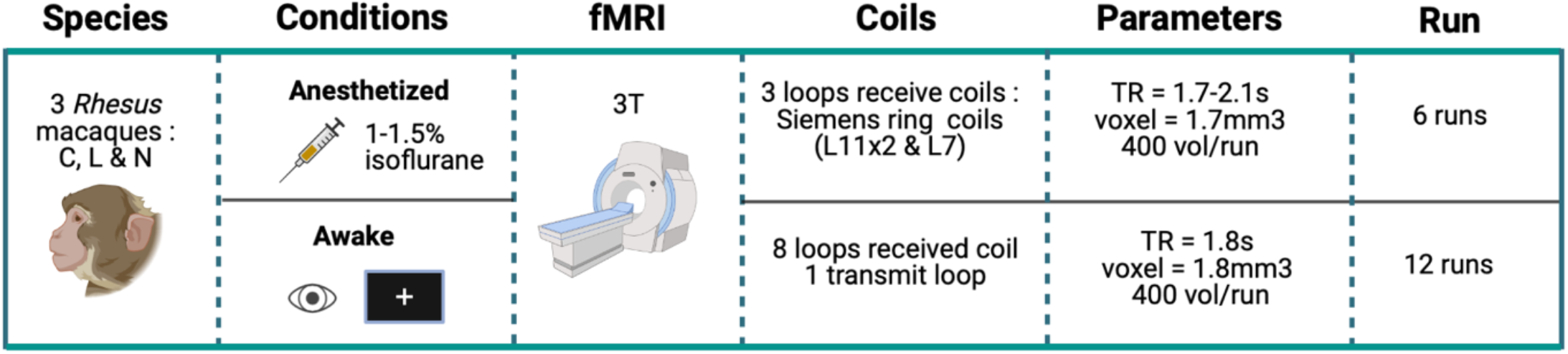
Overview of the experimental design.

| Species | Conditions | fMRI | Coils | Parameters | Run |
| --- | --- | --- | --- | --- | --- |
| 3 *Rhesus* macaques :<br>C, L & N | **Anesthetized**<br>1-1.5% isoflurane | 3T | 3 loops receive coils :<br>Siemens ring coils (L11x2 & L7) | TR = 1.7-2.1s<br>voxel = 1.7mm<sup>3</sup><br>400 vol/run | 6 runs |
|  | **Awake** |  | 8 loops received coil<br>1 transmit loop | TR = 1.8s<br>voxel = 1.8mm<sup>3</sup><br>400 vol/run | 12 runs |

### Anaesthesia impacts the correlation strength between cingulate seeds and lateral frontal ROIs

For each hemisphere of each animal and in each state, we computed the correlation coefficients between the mean signal of cingulate seeds and ROIs in the prefrontal cortex and the motor cortex (Figures 1.A and 2.A). In Figures 1.B and 2.B, the results are displayed on boxplots with each seed represented on the horizontal axis and correlation strength (Z-value) on the vertical axis. In Figures 1.C and 2.C, the plots represent the correlation strength linear trend between each seed and ROIs numbered and ranked from 1 to 7 based on their rostro-to-caudal position (renamed ROIline). The ranking was based on their averaged Y coordinate values across macaque brains and recoded into a numeric axis variable (ROIline): 1) Area 10 (mean Y value across both hemispheres of the 3 macaques = 42, most anterior), 2) Area 46 (mean Y value = 35), 3) Area 9/46 (mean Y value =28.3), 4) Area 44 (mean Y value = 27.5), 5) FEF (mean Y value = 24.2), 6) M1Face (mean Y value = 16.5), 7) M1Hand (mean Y value = 9.2, most posterior). Statistical analysis show that z-values were higher in the awake *versus* anaesthetized runs (main effects of STATE in the left hemisphere: F=28.53, p<9.274e-08; in the right hemisphere: F=14.182, p<0.0002). In both the right (Figure 1) and the left hemispheres (Figure 2), the interaction between the STATE (anaesthetized/awake), the SEED identity (CgS1 to CgS11), and the ROIline (1 to 7) was statistically significant (left hemisphere: df = 10, F=112.683, p<2.2e-16, right hemisphere: df = 10, F=155.259, p<2.2e-16, GLMM with 3 factors (ROIline, SEED, STATE) and 1 random factor (Macaque ID). Note that we observed a main effect of SEED and of ROIline, both in the left (SEED effect: F=330.681, p<2.2e-16; ROIline effect: F=294.134, p<2.2e-16) and the right hemispheres (SEED effect: F=376.094, p<2.2e-16; ROIline effect: F=405.607, p<2.2e-16). When considering hemispheres as an additional factor, the results show a significant interaction between STATE (anaesthetized, awake), SEED (CgS1 to CgS11), ROIline (1 to 8) and HEMISPHERE (left/right): GLMM, fixed effects = hemisphere (left, right), seeds, ROIline, and state, random effect = macaque ID), F=3.88, p<2.757e-05. However, given the small sample size and the lack of a priori hypothesis regarding lateralization, we refrain from discussing further this result.

**Figure 1.**
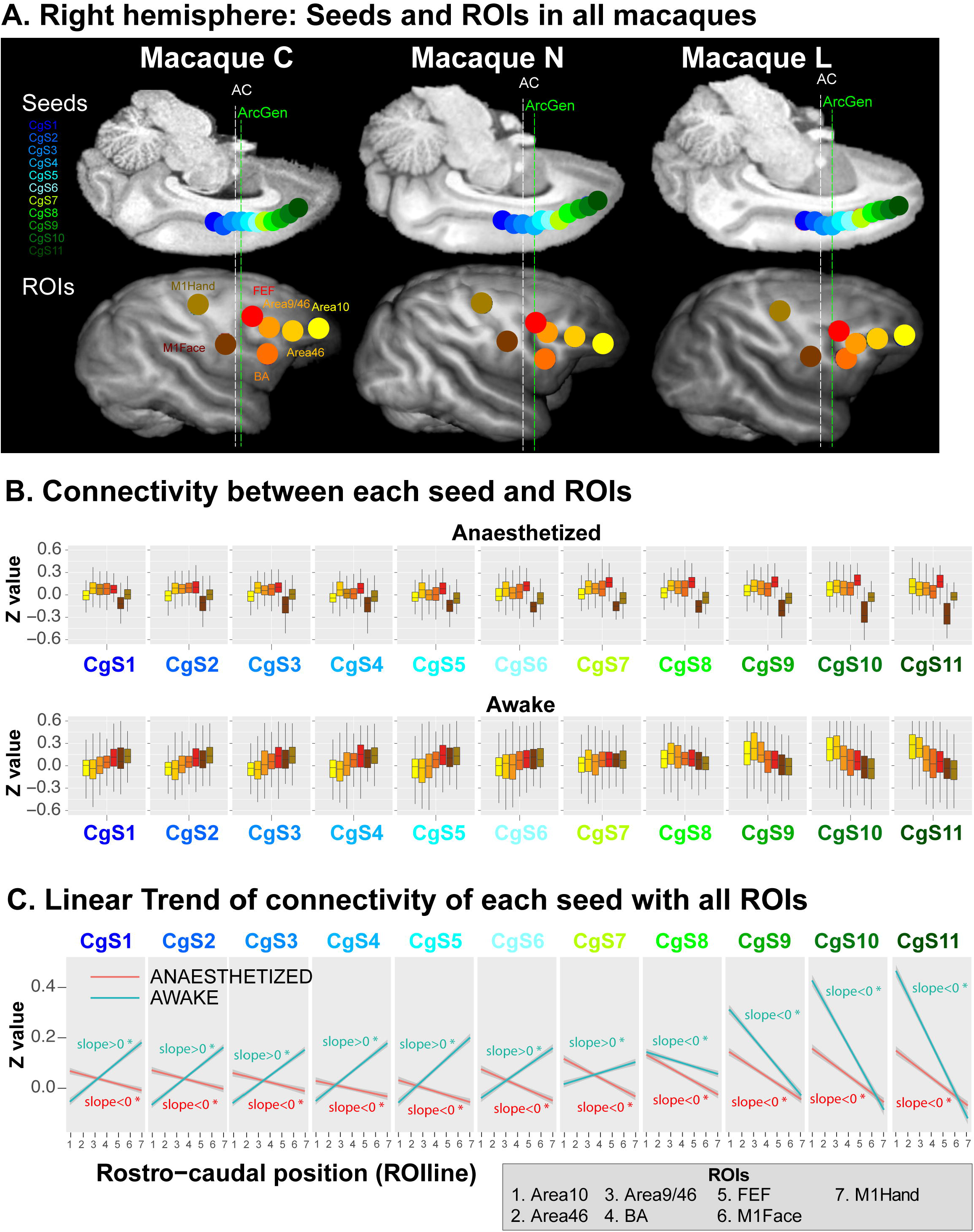
Anaesthesia altered the functional gradient inversion between cingulate regions and lateral prefrontal and motor regions in the right hemisphere. **A.** Position of the seeds and ROIs for the 3 macaques in the right hemisphere. Top part represents the 11 seeds along the cingulate sulcus in an inverse midsagittal section and the bottom part the 7 ROIs in the lateral surface of the brain. The colour conventions are maintained through the rest of the figure. **B**. Boxplots represent the FC between each of the 11 seeds and the ROIs, at the top the anaesthetized condition and at the bottom the awake condition. In the anaesthetized state, seeds CgS1 to CgS11 show stronger connectivity with rostral lateral prefrontal seeds and weaker connectivity with posterior frontal motor ROIs whereas in awake state there is a gradient inversion at CgS7/CgS8 level where caudal seeds CgS8 to CgS11 are stronger connected to caudal frontal motor regions and weaker with rostral lateral prefrontal regions **C**. Linear trend of connectivity between each seed and ROIline in both conditions (ROIs ranked based on their rostro-to-caudal position). No inversion gradient in the anaesthetized state compared to the awake state at the CgS7 and CgS8 transition (GLMM: STATE effect: F=12.469, p<0.0004142, SEED effect: F=380.666, p<2.2e-16; ROIline effect: F=410.959, p<2.2e-16, and effect of STATE*SEED*ROILine interactions df = 10, F=157.147, p<2.2e-16).

**Figure 2.**
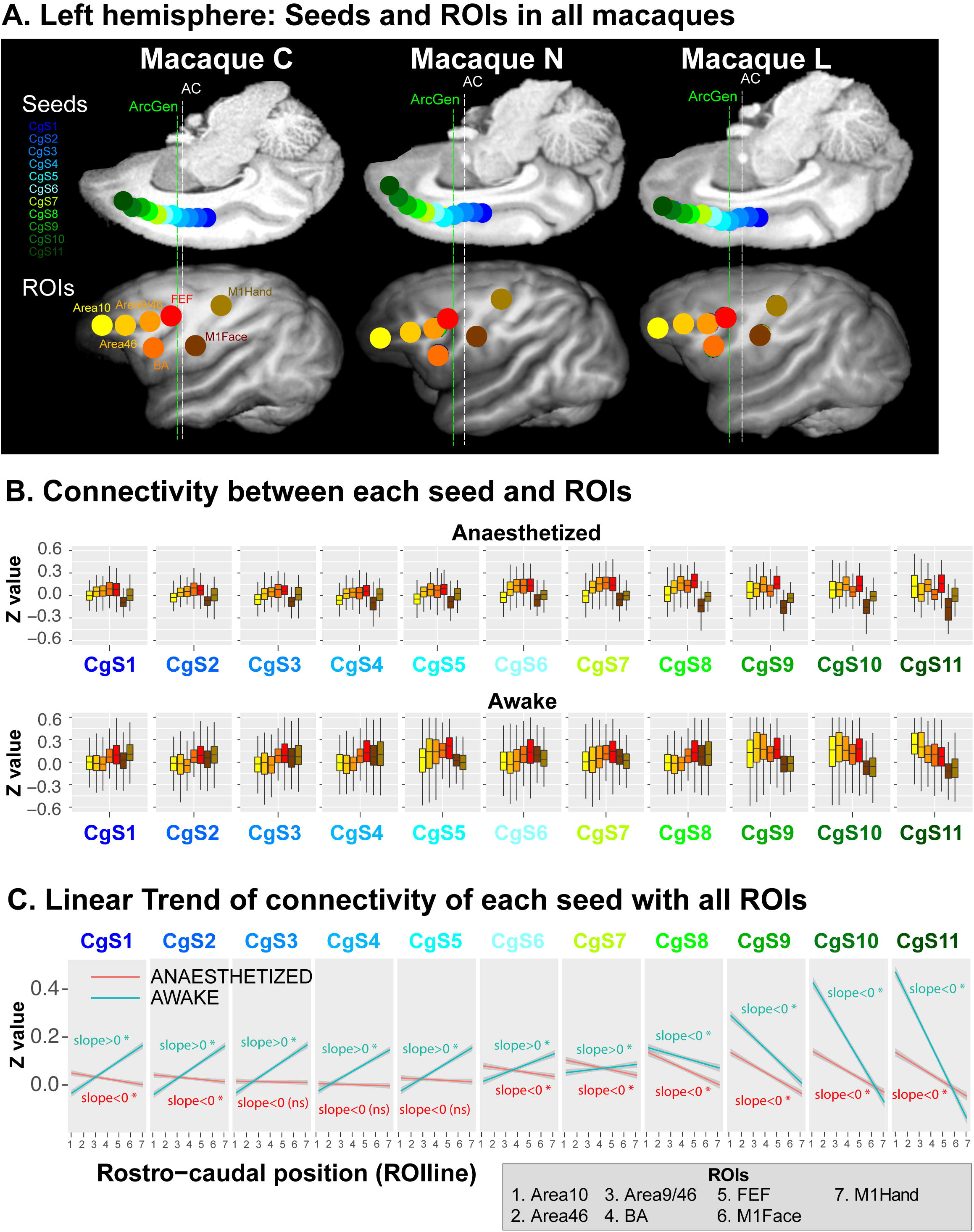
Anaesthesia altered the functional gradient inversion between cingulate regions and lateral prefrontal and motor regions in the left hemisphere. **A.** Position of the seeds and ROIs for the 3 macaques in the left hemisphere. Top part represents the 11 seeds along the cingulate sulcus in an inverse midsagittal section and the bottom part the 7 ROIs in the lateral surface of the brain. **B**. Boxplots represent the FC between each of the 11 seeds and the ROIs, at the top the anaesthetized condition and at the bottom the awake condition. In the anaesthetized state, seeds CgS1 to CgS11 show stronger connectivity with rostral lateral prefrontal seeds and weaker connectivity with posterior frontal motor ROIs whereas in awake state there is gradient inversion at CgS7/CgS8 level where caudal seeds CgS8 to CgS11 are stronger connected to caudal frontal motor regions and weaker with rostral lateral prefrontal regions **C**. Linear trend of connectivity strength between each seed and ROIline (ROIs ranked based on their rostro-to-caudal position) in both conditions. No inversion gradient in the anaesthetized state compared to the awake state at the CgS7 and CgS8 transition (GLMM: STATE effect: F=28.84, p<7.901e-08; SEED effect: F=330.973, p<2.2e-16; ROIline effect: F=294.497, p<2.2e-16, and effect of STATE*SEED*ROILine interactions df = 10, F=112.782, p<2.2e-16)

In the awake state, results show that, in both hemispheres, caudal cingulate seeds CgS1 to CgS7 display stronger connectivity strength with caudal frontal motor ROIs and weaker connectivity strength with rostral prefrontal cortical seeds. Conversely, the rostral cingulate seeds CgS8 to CgS11 display stronger connectivity strength with rostral lateral prefrontal cortical seeds and weaker connectivity strength with caudal frontal motor ROIs. This gradient inversion at the CgS7/CgS8 transition can be visually appreciated in both hemispheres on the boxplots of Figures 1.B and Figure 2.B and the slopes of the linear trends in the correlation strength for each cingulate seed with the rostro-caudal lateral frontal ROIs (Figures 1.C and Figure 2.C, see also Supplemental Table 1 which displays the values of the slopes and corresponding p-values for each seed).

By contrast, in the anaesthetized state, in both hemispheres, all cingulate seeds CgS1 to CgS11, regardless of their position, display stronger correlation strength with rostral lateral prefrontal cortical seeds and weaker correlation strength with caudal frontal motor ROIs (see Boxplots in Figures 1B and 2B and linear trend of connectivity between each seed and the set of ROIs in Figures 1C and 2C). Contrasting with the awake condition, there was no apparent gradient inversion along the axis in the anaesthetized condition.

To better identify the impact of the state (anaesthetized/awake) on the connectivity between each ROIs and all seeds, we performed GLMM with 2 fixed factors (SEED, STATE) and 1 random factor (Macaque ID). The interaction between State and Seed was significant for all ROIs in both the right and left hemispheres (RIGHT: Area 10: df=10, F=49.54 p<2.2e-16; Area46: df=10, F=110.11, p<2.2e-16; Area9/46: df=10, F=9.6, p<5.044e-16; BA: df=10, F=7.26, p<1.6e-11; FEF: df=10, F=14.27, p<2.2e-16; M1Face: df=10, F=2.96, p<0.001; M1Hand: df=10, F=39.724, p<2.2e-16; LEFT: Area 10: df=10, F=21.15, p<2.2e-16; Area46: df=10, F=74.02, p<2.2e-16; Area9/46: df=10, F= 30.53, p<2e-16; BA: df=10, F=6.2, p<1.713e-09; FEF: df=10, F=5.13, p<1.66e-07; M1Face: df=10, F=8.11, p<0.78e-13; M1Hand: df=10, F=30.011, p<2.2e-16). Post-hoc Tukey comparisons revealed that the increased correlation strength between the most caudal cingulate seeds and the motor cortical areas M1Face and M1Hand, and between the most rostral cingulate seeds and lateral prefrontal cortical areas 10, 46, and 9/46, were only present in awake condition, in both hemispheres (Left hemisphere: Figure S4, Right hemisphere: Figure S5). These results suggest that the rostro-to-caudal inversion of gradient was present only in the awake state.

The inversion of gradient occurred on average, across the 3 macaques, at the CgS7/CgS8 transition (i.e. at 10mm anterior to the anterior commissure and on average at 9mm anterior to the genu of the arcuate sulcus -ArcGen-). Note that the transition was observed at slightly different levels for monkeys C, N and L in the two hemispheres (see figures S1, S2 and S3). So, this inversion occurs in a cingulate zone ranging from Y=3.5 to Y=10.5mm from ArcGen.

### Anaesthesia impacts the network connectivity pattern between cingulate seeds and lateral frontal ROIs

To identify the effect of the state (anaesthesia /awake) across the medio-lateral frontal networks, we computed a correlation matrix between all regions defined as seeds and ROIs (i.e. SEED-ROI Z-correlations; SEED-SEED Z-correlations and ROI-ROI Z-correlations; see Materials and Methods, part Statistical Analysis, *Correlations within the frontal cortex and hierarchical clustering)*. The correlation heatmaps are displayed in Figures 3A and 3B for the anaesthetized and awake states, respectively.

**Figure 3.**
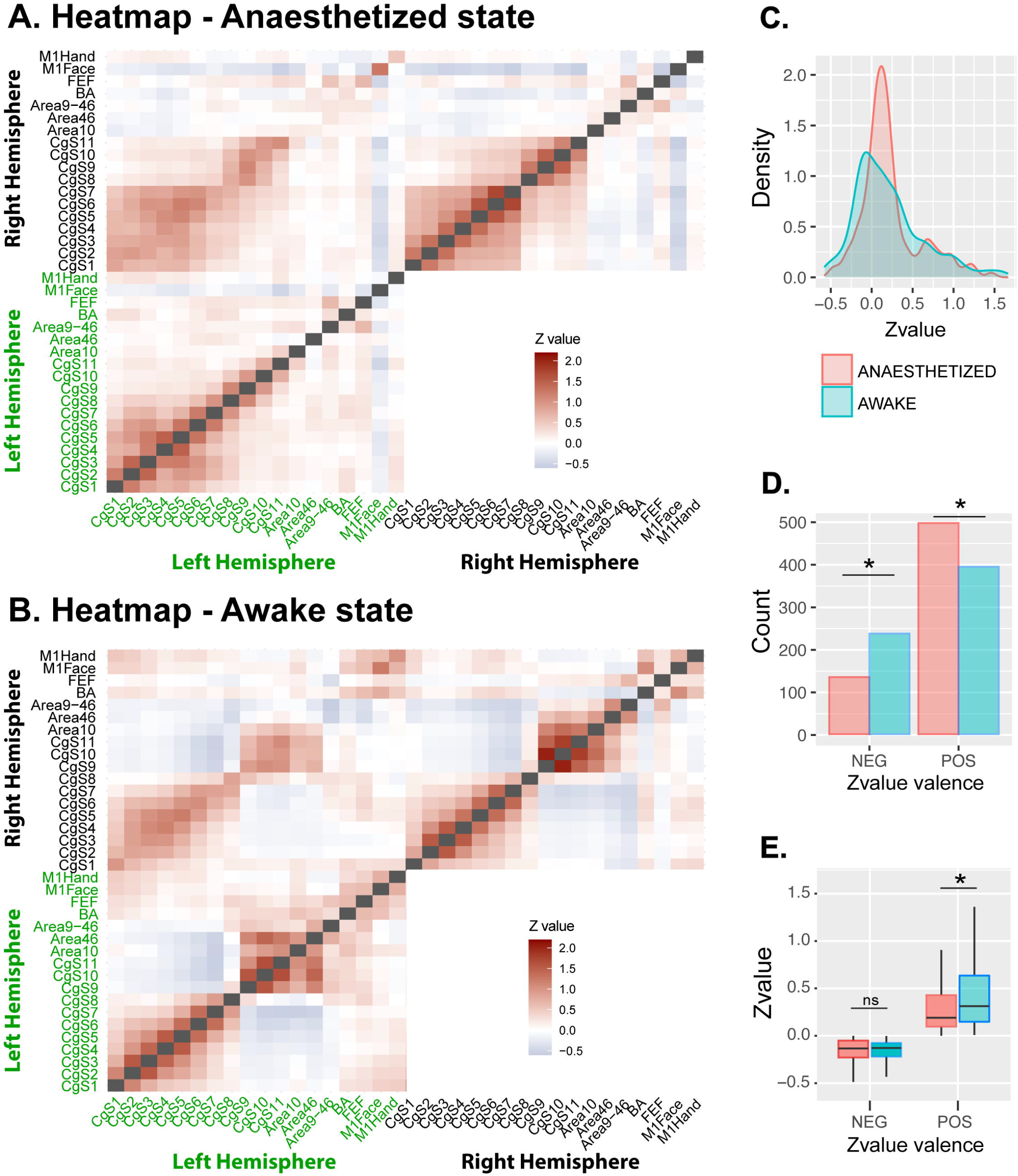
Anaesthesia impacts the Medio-lateral frontal network global network connectivity pattern. **A.B.** Heatmaps of the functional correlations (Z-value) between all seeds and ROIs uni- and bilaterally in the anaesthetized (A) and awake (B) state. Color gradient from red to blue, with red representing positive Z-values and blue negative Z-values. Negative and positive Z-values are differently distributed in awake vs anaesthetized state. The grey diagonal represents auto-correlations. **C**. Density plot representing the distribution of Z-value in both conditions: anaesthetized (in pink) peaked in negative Z-value and awake (in blue) in positive Z-values. **D**. Number of positive and negative Z-values: more negative Z-values in awake vs anaesthetized (26.2% *versus* 39.8%, respectively, Chi2= 35.145, p<3.1e-9) and more positive Z-values in the anaesthetized than awake (73.8% *versus* 60.2%, respectively, Chi2= 17.323, p<3.2e-5). **E**. Boxplots representing the mean Z-values in both positive and negative in each condition. Positive Z-values mean is increased in awake state (estimate = 0.047, p<0.0001, post-hoc Tukey) and no difference on mean negative Z-value (estimate = -0.001, p=0.92, post-hoc Tukey).

Visual inspection of the heatmaps suggest that the pattern of connectivity observed unilaterally is similar to that observed bilaterally, and that negative and positive correlations are not distributed in a similar fashion in anaesthetized and awake states. We further analyzed the distribution of the correlation coefficients (Z-values, positive or negative) within these heatmaps. First, results revealed that the distribution of the Z-values were narrower and the peak of distribution was shifted towards positive values in anaesthetized compared to awake state (peaks at -0.07 and 0.12 in awake and anaesthetized states, respectively, see also histogram of Z-values density in anaesthetized *versus* awake states, Figure 3C). In other words, there were more negative correlations in the awake compared to the anaesthetized state (37.5% *versus* 21.3%, respectively, Chi2= 27.57, p<1.5e-7, proportion test), and conversely more positive correlations in the anaesthetized compared to the awake state (78.7% *versus* 62.5%, respectively, Chi2= 11.703, p<6.2e-4, proportion test, Figure 3D). Second, we found an interaction between the state (anaesthetized/awake) and the sign of the correlation values (positive/negative) on Z-values (df=1, F=9.96, p<0.002, GLM, fixed effect = Z-value valence and STATE, Figure 3E). Although the mean negative Z-values across these networks were not impacted by the state (estimate = -0.005, p=0.75, post-hoc Tukey), positive Z-values were higher in the awake than in the anaesthetized state (estimate = 0.055, p<0.0001, post-hoc Tukey).

### Cingulo-lateral frontal networks are differently organized in the awake *versus* the anaesthetized state

To assess whether the state (anaesthetized/awake) had an impact on the FC in the cingulo-lateral frontal networks, uni- and bilaterally, we performed an unsupervised hierarchical clustering of the seeds and ROIs based on their inter-correlations across all macaques in the awake and anaesthetized states (see Materials and Methods). Resulting dendrograms from the seed and ROI clustering are displayed in Figure 4 along with a heatmap reflecting the correlation strengths between each pair of seed-seed, seed-ROI, and ROI-ROI clusters. In the anaesthetized state, this analysis demonstrated two functional networks between regions of the cingulate cortex (in pink, Figure 4A) and a functional network between regions of the lateral frontal cortex (in purple, Figure 4A), suggesting poor functional interplay between these two entities. By contrast, in the awake state, we found three main functional networks : 1) an anterior cingulate-lateral prefrontal network (dark green, Figure 4B) encompassing

**Figure 4.**
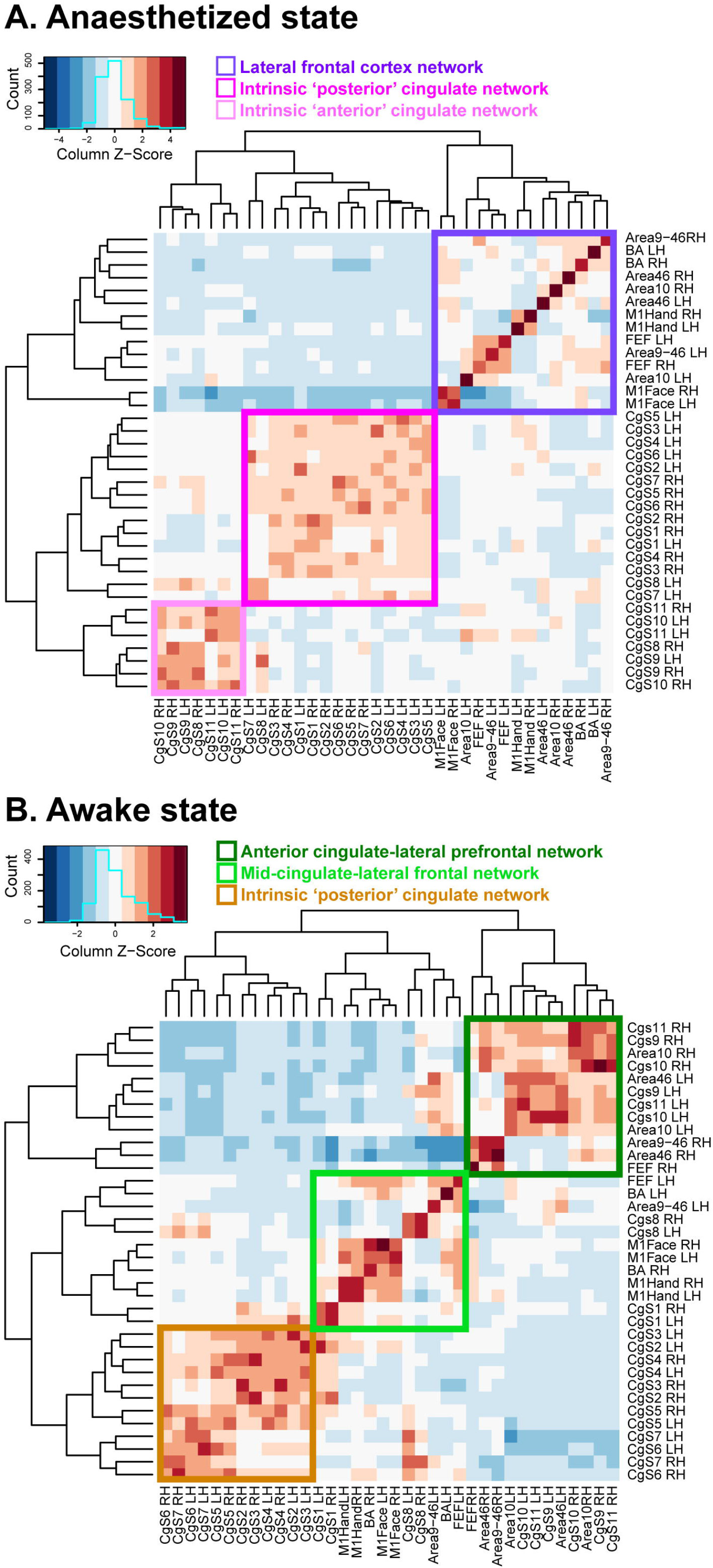
In the large medio-lateral network clustering dissociates three main networks in awake state *versus* two main networks in anaesthetized state. Hierarchical clustering of all seeds and ROIs based on their inter-correlations in anaesthetized (A) and awake (B) state. Dendrograms are represented on the left and top side of the heatmap. **(A)** In the anaesthetized state two main clusters: lateral frontal cortex network (purple) and cingulate network (pink). **(B)** Three clusters are sorted for the awake state: anterior cingulate-lateral network (dark green), posterior cingulate-lateral prefrontal network (light green) and posterior intrinsic cingulate network (light brown).

Area 10, 46, and 9/46, and the three most anterior cingulate areas (CgS9, CgS10, CgS11), 2) a posterior cingulate-lateral prefrontal network (light green, Figure 4B) including Area 9/46, BA, FEF, M1Face and M1Hand, and CgS8 (where the gradient inversion is observed, Figures 1 and 2), and a posterior intrinsic cingulate network (in brown, Figure 4B) comprising cingulate seeds from Cgs1 to CgS7. Note that all these networks are uni-and bilaterally organized. In sum, the analysis provides evidence of a different functional interactions in the cingulo-frontal lateral networks between the awake and anaesthetized states.

### rs-fMRI signal quality check: motion estimation and temporal signal to noise ratio

We assessed the quality of our two datasets to rule out potential confounding effects. We first extracted mean signal from Seeds and ROI regions and then computed for each run and each monkey the temporal signal to noise ratio (tSNR; see Material and Methods). The tSNR was higher in anaesthetized compared to awake sessions (mixed general linear model -GLMM-, fixed effect = state (anaesthetized, awake), random effect (macaque ID), df = 8.654e+05, t-value = -535.16, p<2e-16, Figure 5A). For each of our predefined seeds and ROIs, the tSNR was also higher in anaesthetized compared to awake sessions both in the left and right hemispheres (LEFT: GLMM, fixed effects = state [anaesthetized/awake] and Seeds/ROIs [CgS1 to CgS11, Area 10, Area 46, Area 9/46, BA, FEF, M1Face, M1Hand], random effect [macaque ID], df = 17, F=44.42, p<2.2e-16; RIGHT: df = 17, F=59.03, p<2.2e-16, GLM, Figure 5B).

**Figure 5.**
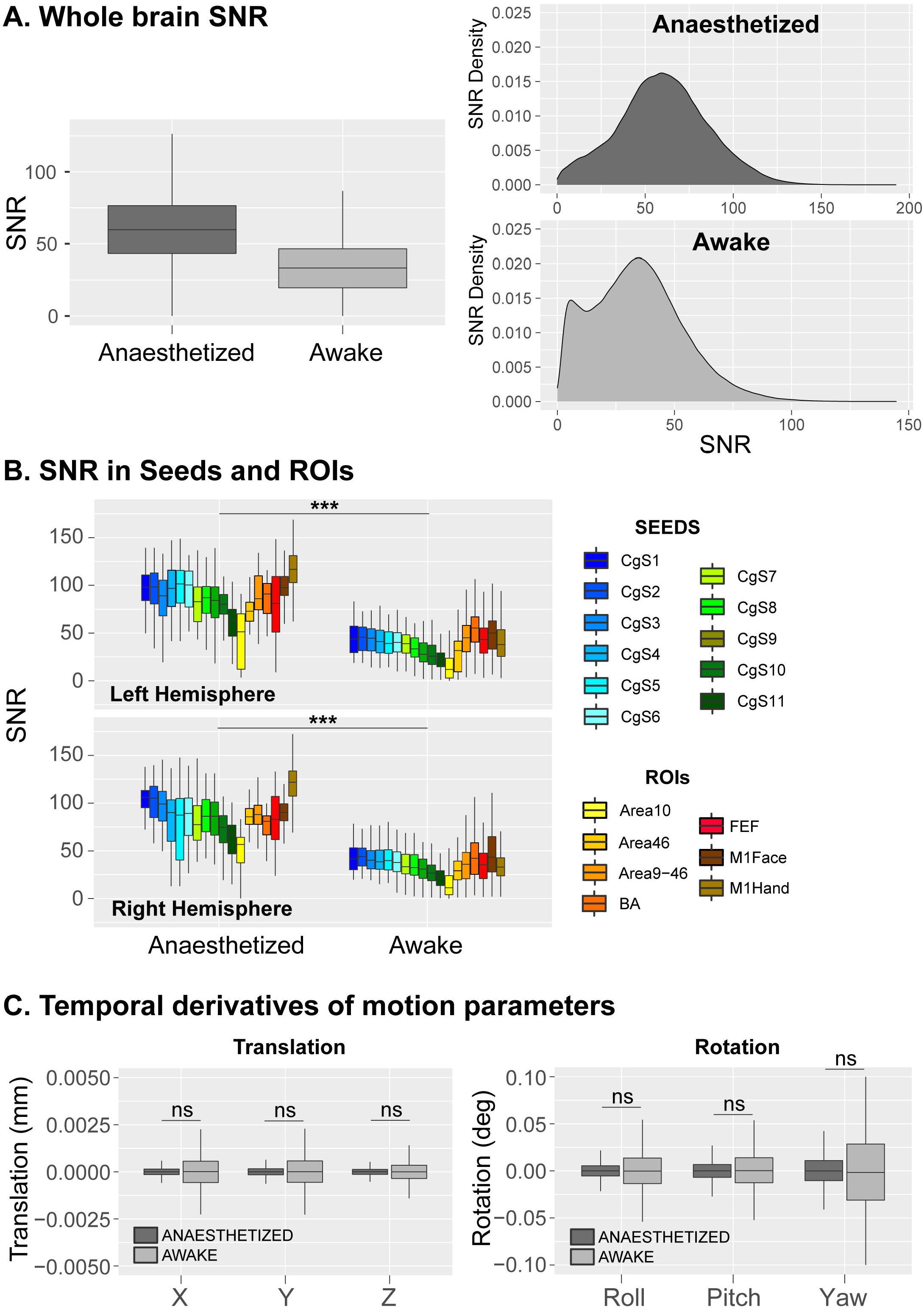
Temporal Signal/Noise ratio and estimated brain movement in the awake and anaesthetized states. **A.** Whole brain temporal signal-to-noise ratio (SNR). Right panel: boxplot representing the tSNR mean in both conditions. GLMM (df = 8.654e+05, t-value = -535.16, p<2e-16) shows a greater tSNR in the anaesthetized compared to awake state. Left panel represents the distribution of tSNR values in the whole brain. **B**. Temporal SNR in each seed and ROI in both hemispheres and conditions: GLMM (df = 17, F=44.42, p<2.2e-16) shows a greater tSNR in the anaesthetized compared to awake state. **C**. Boxplot represents the temporal derivatives of estimated brain movements (translation and rotation), with no significant effect of movement across conditions (GLMM, f = 5, F=0.0049, p>0.05).

The difference in tSNR is reasonably related to the different coils used in the awake *versus* anaesthetized states due to technical limitations (see Material and Methods). Yet, importantly, these differences in tSNR between both states cannot be attributed to variations in estimated brain movements (GLMM, fixed effects = STATE (anaesthetized/awake) or movement types (translations in X, Y, and Z directions; yaw, roll, and pitch rotations), random effect (macaque ID), df = 5, F=0.0049, ns, Figure 5C). Importantly, all results presented above, obtained in awake macaques are similar to those obtained in awake humans with a different coil.

## Discussion

In the present study, we investigated the FC organization between the cingulate cortex and the lateral frontal cortex in monkeys under two states, awake and anaesthetized, using rs-fMRI. In the awake state, we consistently observed for the 3 monkeys a FC organization that follows a rostro-caudal functional gradient such that rostral cingulate regions were more functionally correlated with rostral lateral prefrontal regions (i.e. areas 10, 46 and 9/46) compared to fronto-lateral motor regions. Inversely, caudal cingulate regions were more functionally correlated with frontal lateral motor regions (i.e. M1Face and M1Hand) compared to rostro-lateral prefrontal regions. This FC organization is similar to that previously described in humans (Loh et al., 2018). More importantly, our results reveal that, in the anaesthetized state 1) this functional gradient inversion was not apparent, and 2) the number of negative correlations and the strength of positive correlations within the different parts of the networks were reduced compared to the awake state. These results provide evidence that anaesthesia impacts the FC between the cingulate cortex and the lateral frontal cortex.

### Similar rostro-caudal functional gradient in cingulo-frontal lateral networks in awake humans and monkeys

As discussed above, Loh et al. (2018) recently reported a particular rostro-caudal FC organization within the cingulo-frontal lateral network using rs-fMRI in awake humans. Anterior cingulate regions displayed stronger positive correlations with rostro-lateral prefrontal regions and weaker ones with the lateral caudal motor regions whereas more caudal cingulate regions displayed the reverse pattern. Interestingly, the present study reveals a similar FC organization of the cingulo-frontal lateral network when rhesus macaques were awake and an altered organization when they were anaesthetized.

While in the study of Loh et al. (2018), seeds in the cingulate cortex regions and ROIs in the motor cortical areas were defined based on activation peaks from a fMRI motor mapping task and ROIs in the lateral prefrontal cortex were defined based on local anatomy, all the seeds and the ROIs in the monkey of the present study were identified based on local anatomy. Specifically, based on a previous metanalysis suggesting that the face motor area of CMAr is located about 10mm anterior to the genu of the arcuate (Procyk et al. 2016), we anticipated that this subdivision would roughly correspond to CgS8. Interestingly, the FC profile of this region and that of more anterior cingulate seeds CgS8 to CgS11) that we uncovered in the present study display a similar pattern than that found previously in human and chimpanzee, RCZa (homologue of the macaque CMAr). In the present study and as previously reported in human and chimpanzee (Loh et al. 2018; Amiez et al. 2021), these seed regions display functional coupling with BA, FEF, M1Face, M1Hand in awake state (Figure S4 and S5). Altogether, these data suggest that our seed CgS8 might represent the caudal limit of the homologous of RCZa in human, i.e. CMAr.

Concerning CMAc, it is subdivided into CMAv and CMAd (Dum & Strick, 2002), and these regions are thought to correspond to human RCZp and CCZ (Amiez and Petrides 2014; Loh et al. 2018). We have shown that, in humans, the gradient of connectivity of RCZp and CCZ with the same set of ROIs in frontal cortex displays the reverse pattern of RCZa, i.e. increased connectivity with caudal motor lateral frontal cortical seeds and decreased connectivity with rostral lateral prefrontal cortical seeds. Our results show that this reversed gradient in the awake macaque can be observed from the most caudal seed (CgS1) and CgS7 (+10mm from AC, +8mm from ArcGen, see Figures 1 and 2). We can therefore reasonably conclude that the macaque homologues of the human RCZp and CCZ are both located within these caudal cingulate seeds (CgS1 to CgS7), but that we cannot identify their respective location on the sole basis of our ROI analysis. Importantly, in humans, we were also unable to dissociate RCZp and CCZ on the basis of their correlation profile using the same chosen set of ROIs (Loh et al. 2018).

In sum, based on their FC signatures with the lateral prefrontal and motor cortex in rs-fMRI, we could identify homologies between humans and monkey cingulate motor regions but only when considering data from the awake state. Note that the FC of the rostral cingulate seeds displays similarities between human awake and chimpanzees anaesthetized with propofol (Amiez et al., 2021). As we did not investigate the FC profile for caudal cingulate seeds, we do not know whether a rostro-caudal gradient inversion is also present in chimpanzees. However, it is likely that, as propofol also strongly alters FC (Barttfelda et al., 2015; Uhrig et al., 2018), the resulting gradient would be abolished but future studies should shed light on this matter. In other words, the state – awake vs. anaesthetized - matters when comparing FC patterns using rs-fMRI across species, and the present study shows that a careful attention should be given to interpreting FC patterns in particular of motor and premotor cortex collected under anaesthesia.

### Anaesthesia alters rs-fMRI-identified FC pattern in cingulo-frontal lateral networks

Anaesthesia is characterized by an alteration of the level of consciousness, decreased muscle tone, and altered autonomic responsiveness (Scharf & Kelz, 2013). The degree to which each of these effects is achieved depends both on the anesthetic agent and its dose. Isoflurane is a volatile inhalation anesthetic agent, commonly used in surgical procedures and in-vivo neuroimaging studies in animals, at concentrations varying between 0.75% to 1.5%. In NHP, increased cerebral blood flow (CBF) has been reported with a dose-dependent effect such that higher dose of isoflurane increases CBF due to larger vasodilation (Matta et al., 1999).

Not surprisingly, these changes in CBF induce various alterations of FC in the brain, as measured using rs-fMRI. First, at the whole brain level, the comparison between awake and anaesthetized conditions in both humans and monkeys, suggest that inter-hemispheric correlations exhibited more pronounced reduction compared to intra-hemispheric ones (Hutchison et al. 2014; Hori et al. 2020) and that this effect was dependent on the dose of anesthetics (Hutchison et al. 2014). Second, anaesthesia is also characterized by weaker correlation strength in addition to a decrease in negative correlations or anticorrelations (Barttfeld et al., 2015; Hutchison et al., 2014; Hori et al., 2020; Uhrig et al., 2018; Xu et al., 2019). Our results on the cingulo-frontal lateral networks reveal no difference in inter- or intra-hemispheric correlations. Yet, we found a significant decrease of the positive correlation strength (Z-values, figure 3.E) and a decrease in the number of negative correlations (count, Figure 3.D) in the anaesthetized compared to the awake state. Note that the tSNR significantly differed between the two states, likely because of the different coils used in the two conditions (see methods). However, the decrease in correlation strength under anaesthesia might not result from a lower ability to detect a signal change because the tSNR was higher than in the awake state, and despite the lower tSNR in the awake state, the positive correlations strengths were higher. In addition, it is unlikely that motion, as measured from the estimated brain movements, could explain the differences between the two states as we did not find any significant difference between them. Negative correlations in resting state fMRI studies are highly debated, in particular on whether they artifactually result from data preprocessing strategies (following global signal regression, see for instance Murphy et al., 2009; Saad et al., 2012). In the present study, we did regress out CSF and WM signals, but not the global signal. Therefore, the reduction of negative correlation reported here is a mere feature of the impact of isoflurane on FC patterns in macaque frontal cortex. Importantly, more recent evidence suggests instead that these negative correlations have biological significance. They might highlight regulatory interactions between brain networks and regions (Barttfeld et al. 2015; Uhrig et al., 2018; Kaundinya et al., 2015) and that they may represent a relevant biomarker in some diseases (Ramkiran et al., 2019). Therefore, the larger number of negative correlations found in the awake compared to anaesthetized state might participate in the configuration of the FC signature within the cingulo-frontal lateral networks.

Beyond the general effect of anaesthesia on brain activity at rest, Hutchison *et al*. in 2011, showed in anaesthetized macaques that most of the large-scale resting state networks are topographically and functionally comparable to human ones (Hutchison et al., 2011). However, they also revealed major differences, especially the lack of a dorsomedial PFC component in the default mode network (DMN). The same group subsequently showed that in anaesthetized marmosets, the coactivation strength decreased within large-scale resting-state networks and that the DMN network was particularly impacted, also exhibiting a lack of frontal component compared to the awake state (Hori et al., 2020). On the contrary, in awake NHP, frontal and prefrontal components of the DMN network are not altered (Hori et al., 2020; Mantini et al., 2011). These results underlie the impact of anaesthesia, on large-scale brain functional brain networks and suggest that regions of the frontal cortex might be particularly sensitive to anaesthetic agents. Similarly, our clustering results have shown that the functional interplay between cingulate and lateral frontal cortex can be observed only in the awake state. In the anaesthetized state, only local FC persists as lateral frontal cortical regions and cingulate regions appeared in separate clusters. These results are in line with the hypothesis stipulating that anaesthetics alter cortical activity by biasing spontaneous fluctuations of cortical activity to a more local brain configuration that is highly shaped by brain anatomy (Uhrig et al., 2018). Therefore, anaesthesia might be a potential confound factor that should be considered carefully when comparing FC patterns across species and under different awareness states, especially when considering frontal motor regions (see also Schroeder et al., 2016, Ma et al., 2019). In our study, anaesthesia strongly affects M1 activity and its functional dialogue with the cingulo-prefrontal network (Supplementary figure S4 and S5). This is in line with other findings showing disrupted FC between M1 region and the cingulate cortex, the somatosensory cortex, the FEF with several anesthetic drugs (Uhrig et al., 2018, Schroeder et al. 2016).

### Conclusion

The present study revealed a similar FC signature in cingulo-frontal lateral networks in awake macaque comparable to that previously described in awake human subjects (Loh et al. 2018) using rs-fMRI, suggesting a persevered FC organization of this network from macaque to human. Specifically, rostral seeds in the cingulate sulcus exhibited stronger correlation strength with rostral compared to caudal lateral prefrontal ROIs while caudal seeds in the cingulate sulcus displayed stronger correlation strength with caudal compared to lateral prefrontal ROIs. By comparing this cingulo-frontal lateral network pattern in awake or anaesthetized animals, we found that the inverse rostro-caudal functional gradient was abolished under anaesthesia, suggesting caution when comparing FC patterns across species under different states.

## Materials and methods

### Monkey subjects

For this study, we included 3 monkeys: two females (Monkeys C, 21 years old and N, 9.5 years old) and one male (Monkey L, 9.5 years old) rhesus monkeys (*Macaca mulatta*, 5 - 8 kg). Animals were maintained on a water and food regulation schedule, individually tailored to maintain a stable level of performance for each monkey. All procedures follow the guidelines of European Community on animal care (European Community Council, Directive No. 86–609, November 24, 1986) and were approved by French Animal Experimentation Ethics Committee #42 (CELYNE).

### Surgical procedure

Macaque monkeys were surgically implanted with a PEEK MR-compatible head post (Rogue Research, CA). The surgical procedure was conducted under aseptic conditions. Animals were sedated prior to intubation and they were maintained under gas anaesthesia with a mixture of O2 and air (isoflurane 1-2%). After an incision of the skin along the skull midline, the head fixation device was positioned under stereotaxic guidance on the skull and maintain in place using ceramic sterile screws (Thomas RECORDING products) and acrylic dental cement (Palacos^®^ Bone cements). Throughout the surgery, heart rate, respiration rate, blood pressure, expired CO^2^, and body temperature were continuously monitored. At the completion of the surgery, the wound was closed in anatomical layers. Analgesic and antibiotic treatment were administered for 5 days postoperatively and monkeys recovered for at least 1 month.

### Experimental set up during the awake state

Beginning approximately 4 weeks after the surgery, the monkeys were acclimatized to the head-fixation system and the MRI environment. Monkeys were trained to sit in a sphinx position in an MRI compatible plastic chair (Rogue Research) with their heads fixed, in a mock scanner mimicking the actual MRI environment. During the training, each animal was habituated to view a central cross presented on a screen in front of them. During the scanning sessions, monkeys sat in a sphinx position in the plastic chair positioned within a horizontal magnet (3-T MR scanner; Siemens). Monkeys faced a translucent screen placed 57 cm from their eyes and a white cross (4°x4°) was presented in the center of a black background on the screen at eye level, aligned with their sagittal axis. Eye position was monitored at 1000 Hz during scanning using an eye-tracking system (Eyelink 1000 Plus Long Range). The horizontal (X) and vertical (Y) eye positions from the right eye of each monkey were recorded for each run and each monkey (12 runs for Monkeys C and N and 8 runs for Monkey L as eye movements from 4 runs were not recorded due to technical issues). The calibration procedure involved the central cross and 4 additional crosses (5 degrees of eccentricity), placed in the same plane as the fixation cross. Each point appeared sequentially on the screen and the monkeys were rewarded for orienting and maintaining their gaze towards the cross. The task and all behavioral parameters were controlled by the Presentation® program (Neurobehavioural System). Visual stimulations were projected onto the screen with a projector (CHRISTIE LX501). Monkeys were rewarded with liquid dispensed by a computer-controlled reward delivery system (Crist®) through a plastic tube placed in their mouth. They were rewarded when their eye gaze was within a 4° window around the cross. For each run, we computed the percentage of time the animals spent with their eyes open or fixating. The mean time with eyes open across runs was respectively 69%, 69% and 84% for monkeys L, N and C. Within this time, the percentage of fixation varied from 36-69%, 2-58% and 5-98% for monkeys L, N and C, respectively.

### Rs-fMRI data acquisition

*Table 1* provides an overview of the protocols and scanning parameters under both states, namely awake and anaesthetized (and was generated using BioRender, see BioRender.com). The data from both states were acquired from the same scanner, in a 3T Siemens Magnetom Prisma MRI scanner (Siemens Healthcare, Erlangen, Germany). Due to the specific constraints in the anaesthetized (stereotaxic frame) and awake (MRI chair) states, we could not use the same coil systems in the 2 conditions.

### Anaesthetized state

Prior to anaesthesia, monkeys were injected with glycopyrrolate, an anticholinergic agent that decreases salivary secretion (Robinul; 0.06mg/kg). Twenty minutes after, anaesthesia was induced with an intramuscular injection of tiletamine and zolazepam (Zoletil; 7 mg/kg). The animals were then intubated and ventilated with oxygen enriched air and 1% Isoflurane (for monkeys N and L) or 1.5% Isoflurane (for monkey C) throughout the duration of the scan. An MRI-compatible stereotaxic frame (Kopf, CA, USA) was used to secure the head and reduce variability in the measure. Monkeys were placed in a sphinx position with their head facing the back of the scanner. Breathing volume and frequency were set based on the animal weight. During the scan, physiological parameters including heart rate and ventilation parameters (spO2 and CO2) were monitored. Body temperature was also measured and maintained using warm-air circulating blankets. The anaesthetized resting-state acquisitions were performed about 2 hours after anaesthesia induction and at least 1 hour after first inhalation of isoflurane.

Three received loop coils were used for the acquisition: 2 L11 Siemens ring coils were placed on each side of the monkey’s head and 1 L7 Siemens above the monkey’s head. A high resolution T1-weighted anatomical scan was first acquired for each of the 3 monkeys (MPRAGE, 0.5mm3 isotropic voxels, 144 slices, TR=3000ms, TE=366ms). Resting-state functional images were obtained in an ascending order with a T2*-weighted gradient echo planar images (EPI) sequence with the following parameters: for Monkey L and N, TR=1700ms, TE = 30ms, flip angle = 75°, FOV = 400×300mm, 25 slices, voxel size: 1.6mm^3^. For Monkey C, TR= 2000ms, TE = 30ms, flip angle = 75°, FOV= 480 × 336, 31 slices, voxel size: 1.8mm^3^ for monkey C. We collected 6 runs for monkeys L and C and 5 runs for monkey N with 400 volumes per run (12 min each) for a total of 6800 volumes across the 3 animals. The different runs were acquired during the same session for each animal.

### Awake state

Data were acquired using a custom-made 8 channel receive surface coil positioned around the head, and a circular transmit coil positioned above the head (Mareyam et al., In: Proceedings of the 19th Annual Meeting of ISMRM, Montreal, Canada, 2011). Functional images were acquired in an ascending order with a total of 12 runs of 12 min per monkey (400 volumes/run, corresponding to a total of 4800 volumes per monkey) across different scan sessions. We used a BOLD-sensitive T2* weighted echo planar sequence with the following parameters TR = 1800ms, TE=27ms, flip angle = 75°, FOV = 480×336 mm, voxel size = 1.8 mm isotropic, 30 slices. Throughout the scan duration, macaques were either fixating the central cross presented on the screen or eyes opened for the duration of the run. Each functional imaging acquisition was preceded by a T1-weighted low-resolution anatomical scan with a MPRAGE sequence (TR = 2500ms, TE=2.44ms, flip angle = 8°, FOV = 128×128 mm, voxel size = 0.9 mm isotropic, 64 volumes).

### Rs-fMRI data analysis

The preprocessing of resting-state scans was then performed with SPM 12. The first 5 volumes of each run were removed to allow for T1 equilibrium effects. First, we performed a slice timing correction using the time center of the volume as reference. The head motion correction was then applied using rigid body realignment. Then, images were skull-stripped using the bet tool from the FSL software (https://fsl.fmrib.ox.ac.uk/fsl/fslwiki/BET, Jenkinson et al. 2005). Using the AFNI software (Cox, 1996), the segmentation of each brain of each session (anaesthetized and awake sessions) was performed on skull-stripped brains. To ensure optimized inter-session and inter-subject comparisons, both anatomical and functional images were then registered in a common atlas space, CHARM/SARM (Jung et al. 2020, Reveley et al. 2017, see https://afni.nimh.nih.gov/pub/dist/doc/htmldoc/nonhuman/macaque_tempatl/atlas_charm.html). Temporal filtering was then applied to extract the spontaneous slowly fluctuating brain activity (0.01–0.1Hz). Finally, linear regression was used to remove nuisance variables (the cerebrospinal fluid, white matter signals from the segmentation, and volumes containing artefacts as detected by the ART toolbox, https://www.nitrc.org/projects/artifact_detect/) and spatial smoothing with a 4-mm FWHM Gaussian kernel was applied to the output of the regression.

#### Seed Selection in the MCC

The seeds consisted of 2.5mm radius spheres positioned in the cingulate sulcus (covering both ventral and dorsal banks of the cingulate sulcus) in both hemispheres, starting 10 mm posterior to the anterior commissure to the rostral end of the cingulate sulcus, and spaced from 2.5mm each, for a total of 11 seeds. (CgS1, CgS2, CgS3, CgS4, CgS5, CgS6, CgS7, CgS8, CgS9, CgS10 and CgS11, see Figure 1 for the positioning of seeds in the right hemisphere and Figure 2 for the left hemisphere).

Given that, in macaques, CMAr is located about 10mm anterior to the genu of the arcuate, we anticipated that this subdivision would roughly correspond to CgS8 (Procyk et al. 2016). Caudal seeds CgS1 to CgS7 correspond to Y values -5 to +10 (with AC at Y0) whereas CgS8 to CgS11 correspond to Y values +12.5 to +20. As it can be appreciated in Figures 1 and 2, the genu of the arcuate sulcus (ArcGen) is at the level of CgS4 (i.e. at Y=+2.5). According to our previous meta-analysis aiming at identifying the location of the rostral cingulate motor area (CMAr) in macaques (Procyk et al. 2016), the face motor area CMAr is located about 10mm anterior to ArcGen.

#### Selection of Regions of Interest (ROIs)

For a stricter comparison of the present results with results obtained in our previous studies which assessed the FC of the cingulate sulcus in the human (Loh et al., 2018) and in the chimpanzee (Amiez et al., 2021), we used the same ROIs (see Figure 1 for the positioning of ROIs in the right hemisphere and Figure 2 for the left hemisphere). Each ROI consisted of a sphere with a 2.5mm radius.

### ROI Selection in Motor Cortical Areas

For each subject, 3 ROIs within the motor cortex of both hemispheres were identified based on sulcal morphology. These included the hand motor region –M1Hand–and the primary face motor region within the ventral part of the posterior part of the precentral gyrus –M1Face– (Graziano et al., 2002; He et al., 1993; Luppino, G., & Rizzolatti 2000). We also included the frontal eye field –FEF–, located in the rostral bank of the arcuate sulcus, at the level of the genu of the arcuate sulcus (Bruce et al. 1985, Amiez and Petrides, 2009).

### Selection of ROIs in the prefrontal cortex

For each subject, 4 ROI locations within the left prefrontal cortex were identified based on local anatomy. On a rostro-caudal axis:

- The frontopolar cortex –Area 10–. It occupies the rostral part of the principalis sulcus (see Petrides and Pandya, 1994).
- Area 46 and 9/46 of the dorsolateral prefrontal cortex -DLPFC-. Area 9/46 occupies the caudal part of the sulcus principalis, and area 46 occupies the sulcus principalis between area 9/46 and area 10 (see Petrides and Pandya, 1994).
- Broca’s region: Area 44 has been shown to occupy the fundus of the ventral part of the arcuate sulcus (Petrides et al., 2005).

### Temporal signal to noise ratio (tSNR)

The mean signal from Seeds and ROI regions was extracted using the AFNI software. For each run and each monkey, we computed the temporal SNR (tSNR - average intensity of time series divided by the standard deviation) across the brain, in each seed and in each ROI, in both states, anaesthetized and awake (Figure 5). The tSNR within the whole-brain, within each seed- and each ROI in awake *versus* anaesthetized state was compared using generalized linear mixed-models for each hemisphere separately (package lme4, R statistical software: https://www.r-project.org/) with the anaesthetized/awake state as fixed effect and the macaque identity as a random factor.

### Estimated brain movements

For each session, we also computed the six temporal derivatives of the estimated brain movements (three translations and three rotations) in each session of all monkeys (see Power et al. 2012, see Figure 5). These temporal derivatives in awake *versus* anaesthetized state were compared using generalized linear mixed-models (package lme4, R statistical software: https://www.r-project.org/) with the anaesthetized/awake state as fixed effect and the macaque identity as a random factor.

### Correlations between seeds and ROIs

For each hemisphere of each animal, Pearson correlation coefficients between the seeds with the various ROIs in the prefrontal cortex and the motor cortex were computed and normalized using the Fisher’s r-to-z transform formula. The significant threshold at the individual subject level was set to Z = 0.1 (p < 0.05). These normalized correlation coefficients, which corresponded to the FC strength between each seed and each ROI in individual brains, were subsequently processed with R software for all the following analyses.

To identify the impact of the state (awake or anaesthetized) on the connectivity profile of each seed with the various lateral frontal ROIs, we constructed boxplots representing the correlation strength of each seed location with each of the ROIs in each state and in each hemisphere. Note that given the sample size, and as we did not have any a-priori hypothesis, we refrain from drawing strong conclusions about a lateralization effect within this network and carried out the analysis thereafter from each hemisphere separately. As carried out previously in humans (Loh et al, 2018), we then characterized, in both states, the rostro-caudal functional axis based on the correlation profiles of each seed with the lateral frontal cortex by estimating linear trends in their correlation strength (for details, see Methods in Loh et al., 2018). The 7 ROIs were first ranked along a rostro-caudal axis based on their averaged Y coordinate values across macaque brains and recoded into a numeric axis variable (ROIline): 1) Area 10 (most anterior), 2) Area 46, 3) Area 9/46, 4) Area 44, 5) FEF, 6) M1Face, 7) M1Hand (most posterior). We then performed multiple linear regressions on the correlation z values with seed identity (CgS1 to CgS11), state (awake or under anaesthesia) and the linear axis variable (ROIline) as predictors for each hemisphere separately or considering hemisphere as an additional factor. We assessed whether the linear trends (slopes) observed for each seed were identical or not in each sulcal morphology using generalized linear mixed-models -GLMM- with ROI identity (Area 10, etc.), state (anaesthetized/awake), and the linear axis variable (ROIline, values 1-7) as fixed effect and the macaque identity and run identity as random factors, followed by Posthoc Tukey comparisons (performed with lsmeans package, https://cran.r-project.org/web/packages/lsmeans/lsmeans.pdf).

We also assessed the correlation z values between each ROI and all seeds separately for each hemisphere using a GLMM with seed identity (CgS1 to CgS11) and state (awake or under anaesthesia) as fixed effect and the macaque identity as random factor followed by post-hoc Tukey comparisons.

### Correlations within the frontal cortex and hierarchical clustering

To assess the impact of anaesthesia on the overall functioning of the network composed by the various seeds and ROIs, in each hemisphere, we averaged the normalized correlation coefficients for each seed-seed, seed-ROI, and ROI-ROI pairing across runs and macaques. Correlation heatmaps were then generated in both states, anaesthetized and awake. Note that autocorrelations were not taking into account in the statistical analysis.

Then, unsupervised hierarchical clustering was performed for the 11 seeds within the cingulate sulcus and the 7 ROIs within the lateral frontal cortex. The method is described in Loh et al. (2018). To summarize, this clustering method defines the cluster distance between two clusters to be the maximum distance between their individual components (using the hclust function in R, see http://www.r-tutor.com/gpu-computing/clustering/hierarchical-cluster-analysis). At every stage of the clustering process, the two nearest clusters are merged into a new cluster. The process is repeated until the whole data set is agglomerated into one single cluster. The outcome was used to construct dendrograms and heatmaps. To better display clusters across ROIs, values in the heatmaps were normalized (z-scored) by column. Therefore, values (and sign) in the heatmap do not represent actual connectivity measures.

## Acknowledgements

This work was supported by the French National Research Agency (C. Amiez: ANR-18-CE37-0012-01; F. Hadj-Bouziane: ANR-15-CE37-0003). C.A., F.H.-B., and E.P. are employed by the Centre National de la Recherche Scientifique. C.G., C.A., D.A.-C., E.P., C.W., J.S are supported by the labex CORTEX ANR-11-LABX-0042 of Université de Lyon. We thank Gislène Gardechaux for technical assistance in training awake macaques. We also thank Franck Lamberton for technical help in acquiring neuroimaging data in awake macaques.

