## Supplemental figures and table for "Frontal cortical functional connectivity is impacted by anaesthesia in macaques": Giacometti_Article_supplemental_material.pdf

### Supplementary material

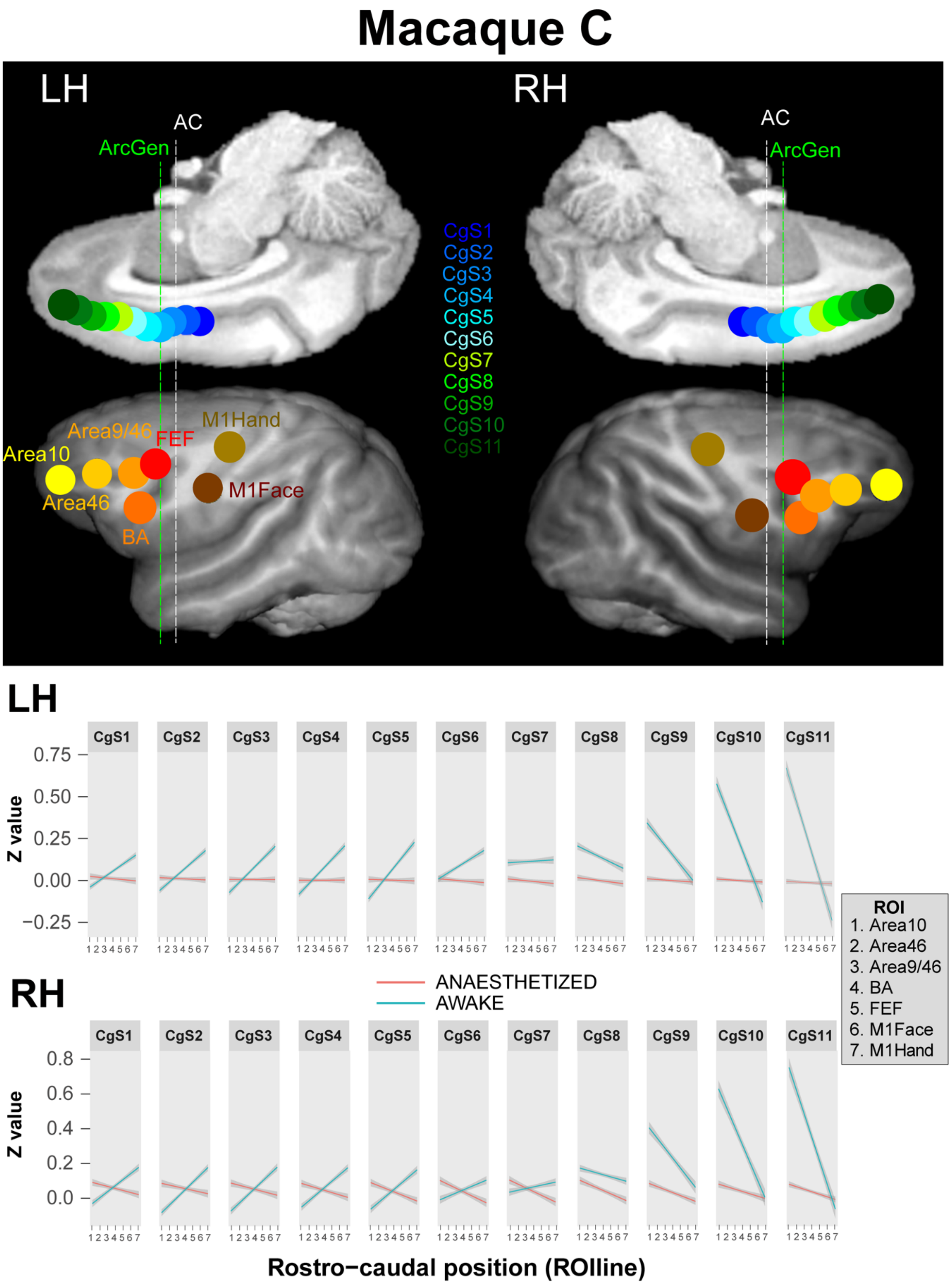


**Figure S1.** Macaque C : alteration of the functional connectivity (FC) gradient inversion between cingulate regions and lateral prefrontal and motor regions in the left and right hemisphere . Top part. Seeds and ROIs localization on the medial and lateral brain surface, respectively, on both the left and the right hemisphere. Bottom part. Plot representing the linear trend of the FC between Seeds and ROIline (ROI ranked by their rostral-to-caudal position) in awake (blue) and anaesthetized (red) state in the right and left hemisphere : for each of the seeds ROIline are represented on the horizontal axis and correlation strength (Z-value) on the vertical axis). In awake state, the inversion of the functional gradient appeared at CgS7/CgS8 transition in both hemispheres whereas this inversion is not apparent in the anaesthetized state. Abbreviation: LH, left hemisphere; RH, right hemisphere.

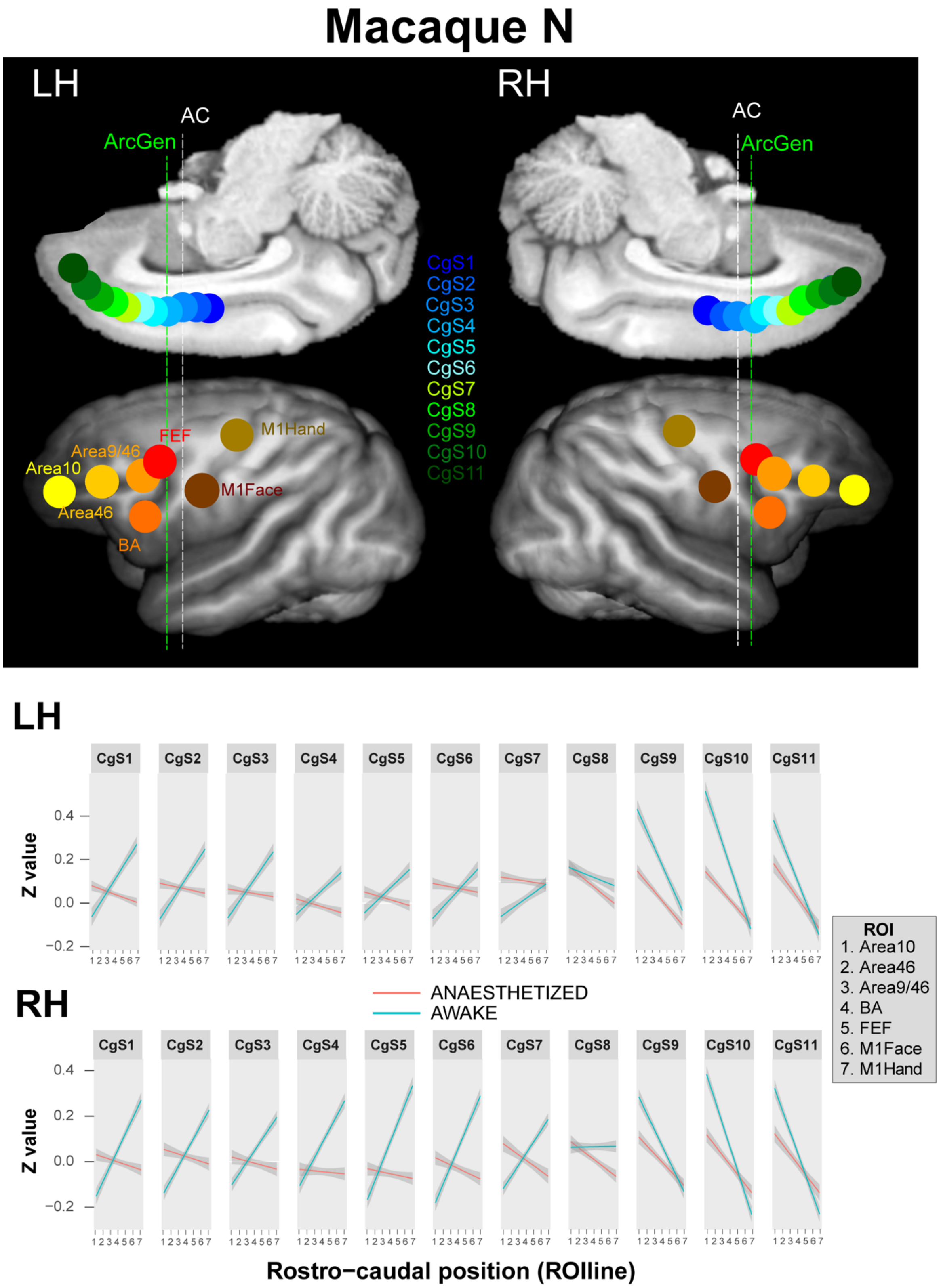


**Figure S2.** Macaque N: alteration of the FC gradient inversion between cingulate regions and lateral prefrontal and motor regions in the left and right hemisphere. The Legend is the same as Macaque C (Figure S1). In awake state, the inversion of the functional gradient appeared at the CgS7/CgS8 transition in the left hemisphere and at the CgS8/CgS9 transition in the right hemisphere whereas this inversion is not apparent in the anaesthetized state.

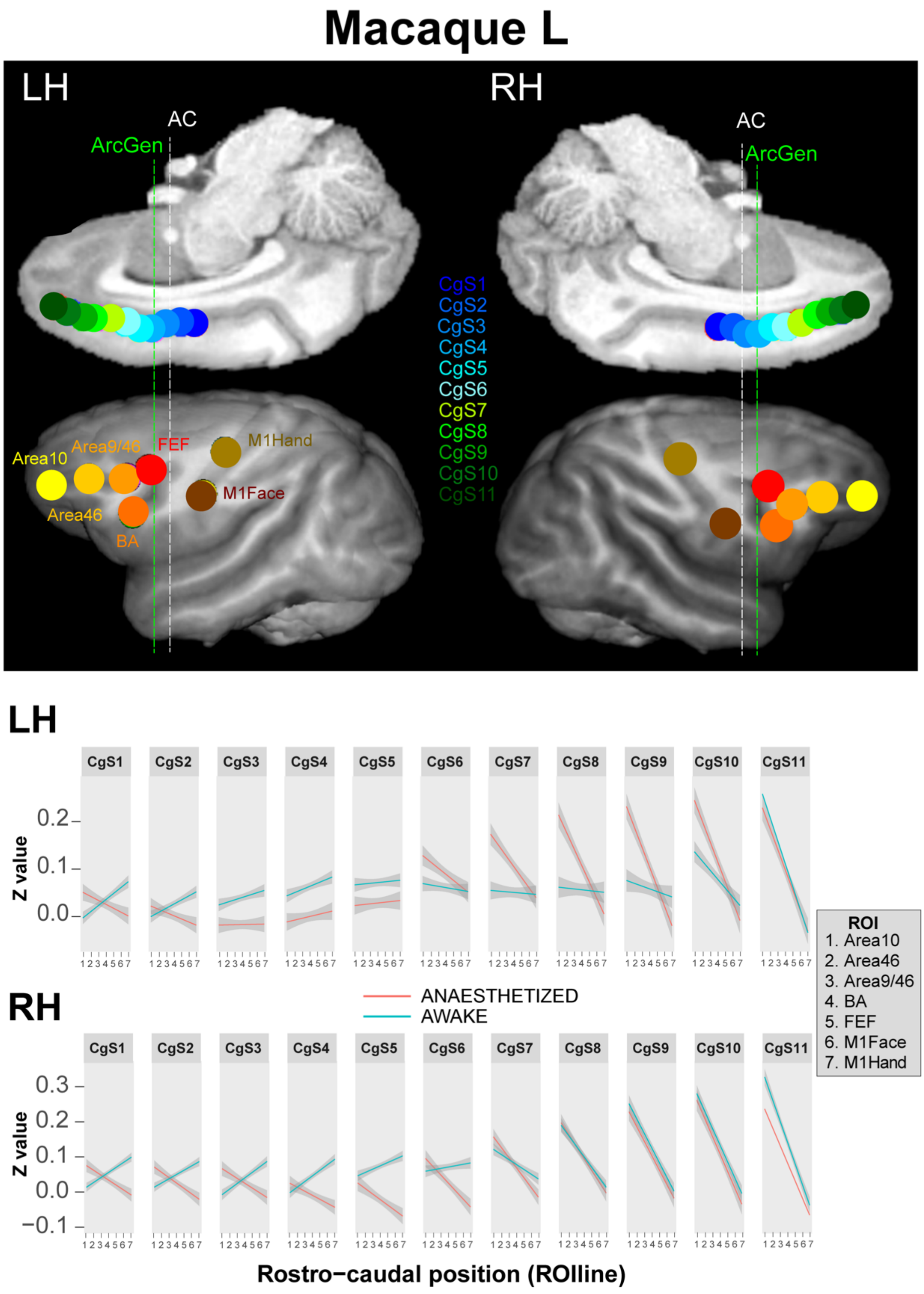


**Figure S3. Macaque L: alteration of the FC gradient inversion between cingulate regions and lateral prefrontal and motor regions in the left and right hemisphere.** The Legend is the same as Macaque C (Figure S1). In awake state, the inversion of the functional gradient appeared at the the CgS5/CgS6 transition in the left hemisphere and at the CgS6/CgS7 transition in the right hemisphere whereas this inversion is not apparent in the anaesthetized state.

#### Right Hemisphere: Intra-hemispheric connectivity between each ROI and all seeds

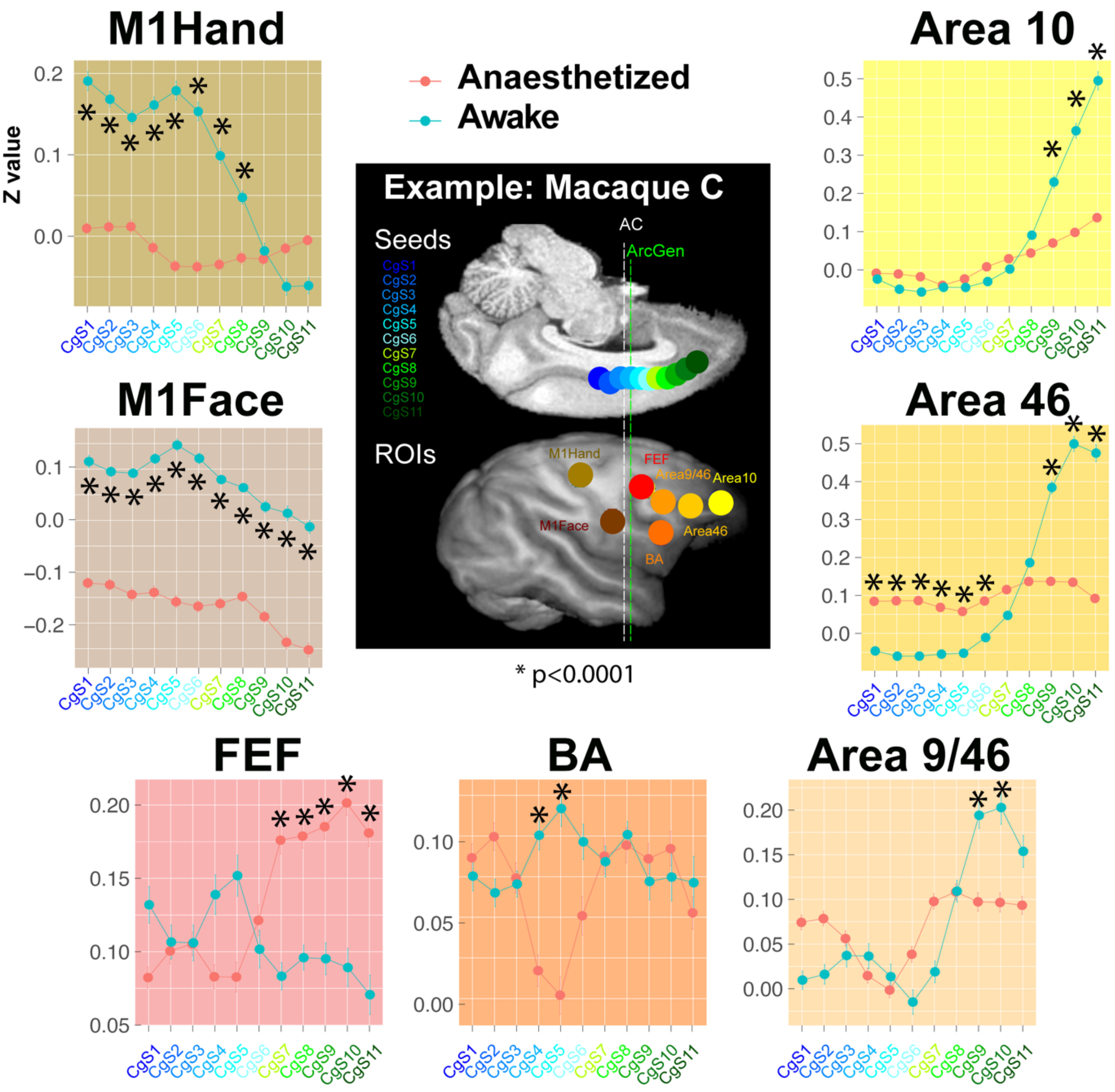


**Figure S4. Intra-hemispheric connectivity between each ROI and all seeds in the right hemisphere**. Seeds and ROIs are displayed on Macaque C' MRI brain surface. The FC between each ROI and all seeds are represented in both awake (blue) and anaesthetized (red) state. (*) represent significant differences in correlations strength between awake and anaesthetized state (Area 10: df=10, F=21.2 p<2.2e-16; Area46: df=10, F=75.1, p<2.2e-16; Area9/46: df=10, F=30.8, p<2e-16; BA: df=10, F=6.2, p<1.6e-9; FEF: df=10, F=5.2, p<1.2e-7; M1Face: df=10, F=8.2, p<2.2e-13; M1Hand: df=10, F=30.7, p<2.2e-16) : the increase in correlation strength between the most caudal cingulate seeds with motor cortical areas M1Face and M1Hand, and between the most rostral cingulate seeds with lateral prefrontal cortical areas 10, 46, and 9/46 is only present for the awake condition.

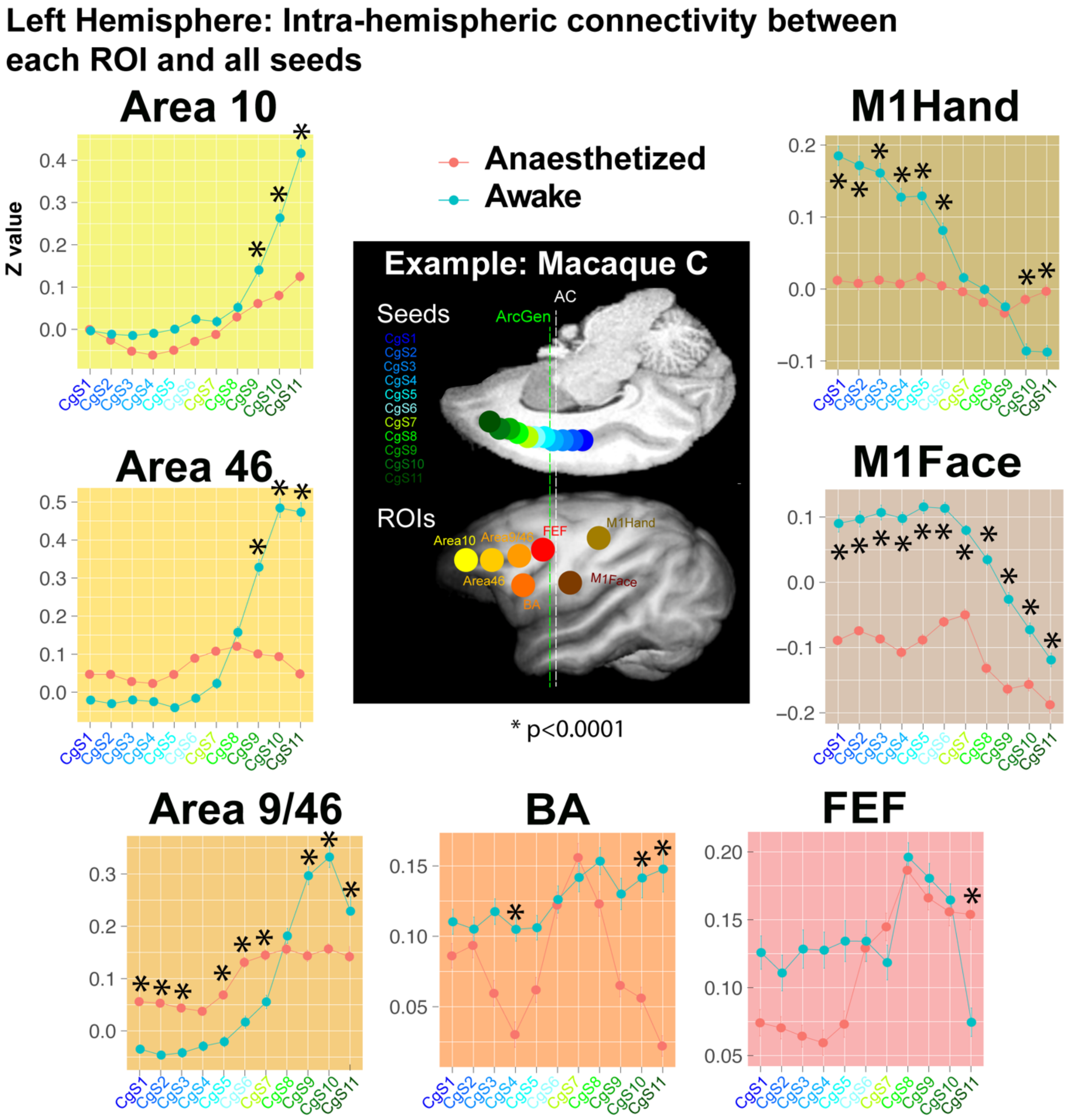


**Figure S5. Intra-hemispheric connectivity between each ROI and all seeds in the left hemisphere**. See Figure S4's legend. (*) represent significant differences in correlations strength between awake and anaesthetized state (Area 10: df=10, F=50.5, p<2.2e-16; Area46: df=10, F=113.2, p<2.2e-16; Area9/46: df=10, F= 9.7, p<2.5e-16; BA: df=10, F=7.3, p<1.4e-11; FEF: df=10, F=14.6, p<2.2e-16; M1Face: df=10, F=3, p<0.0008; M1Hand: df=10, F=40.6, p<2.2e-16). The conclusions are identical than in the rigth hemisphere (see Figure S4).

**Supplemental Table 1. Slopes and p-values of the linear regressions described in Figures 2 and 3.** Significant positive and negative slopes are outlined in orange and blue, respectively.

| | | CgS1 | CgS2 | CgS3 | CgS4 | CgS5 | CgS6 | CgS7 | CgS8 | CgS9 | CgS10 | CgS11 |
|---|---|---|---|---|---|---|---|---|---|---|---|---|
| **Right Hemisphere** | | | | | | | | | | | | |
| AWAKE | Slope | 0.04 | 0.04 | 0.04 | 0.04 | 0.04 | 0.03 | 0.01 | -0.01 | -0.06 | -0.09 | -0.1 |
| | P | 9.7e-83 | 8.3e-93 | 2.2e-80 | 1.4e-77 | 9.4e-77 | 1e-49 | 9.1e-15 | 2.2e-15 | 2.1e-104 | 3.3e-149 | 1.7e-167 |
| ANESTHETIZED | Slope | -0.01 | -0.01 | -0.01 | -0.01 | -0.01 | -0.02 | -0.02 | -0.03 | -0.03 | -0.04 | -0.04 |
| | P | 2.9e-13 | 4.7e-11 | 1.5e-9 | 1.3e-7 | 1.1e-12 | 2.5e-22 | 7.5e-30 | 2.8e-36 | 8e-45 | 9.6e-46 | 1.3e-47 |
| **Left Hemisphere** | | | | | | | | | | | | |
| AWAKE | Slope | 0.03 | 0.03 | 0.03 | 0.03 | 0.03 | 0.02 | 0.006 | -0.01 | -0.05 | -0.08 | -0.1 |
| | P | 7.8e-58 | 3.5e-61 | 1.9e-56 | 9.8e-49 | 1.3e-46 | 2.2e-19 | 0.006 | 2.4e-10 | 0.2e-65 | 4.8e-134 | 6.9e-200 |
| ANESTHETIZED | Slope | -0.008 | -0.005 | -0.00008 | -0.002 | -0.003 | -0.007 | -0.01 | -0.02 | -0.03 | -0.03 | -0.03 |
| | P | 8.6e-7 | 0.003 | 0.58 (ns) | 0.36 (ns) | 0.13 (ns) | 0.0001 | 2.3e-7 | 2.2e-24 | 3.2e-38 | 2.2e-37 | 3.8e-35 |
